# Identification of a new substrate for the ribosome associated endoribonuclease Rae1 reveals a link to the *B. subtilis* response and sensitivity to chloramphenicol

**DOI:** 10.1101/2023.02.16.528812

**Authors:** Valentin Deves, Aude Trinquier, Laetitia Gilet, Jawad Alharake, Magali Leroy, Ciarán Condon, Frédérique Braun

## Abstract

Rae1 is a well-conserved endoribonuclease among Gram-positive bacteria, cyanobacteria and the chloroplasts of higher plants. We have previously shown that Rae1 cleaves the *Bacillus subtilis yrzI* operon mRNA in a translation-dependent manner, within a short open reading frame (ORF) called S1025, encoding a 17-amino acid (aa) peptide of unknown function. Here, we map a new Rae1 cleavage site in the *bmrBCD* operon mRNA encoding a multidrug transporter, within a previously unannotated 26-aa short ORF that we have named *bmrX*. Similar to S1025, Rae1 cleavage within *bmrX* is both translation- and reading frame-dependent. Both mRNAs were previously shown to be induced by the presence of the protein synthesis inhibitor chloramphenicol (Cm). Strikingly, a *rae1* deletion strain shows greater resistance to Cm than the wild-type strain, while its over-expression leads to increased Cm sensitivity, suggesting a link to translation quality control. Consistent with this, we show that cleavage by Rae1 promotes ribosome rescue by the tmRNA. Globally, our data point to a role of Rae1 in mRNA surveillance by eliminating mRNAs that encounter problems with translation.

## INTRODUCTION

In *Bacillus subtilis*, mRNA degradation is typically catalysed either by the RNase J1/J2 complex that degrades RNA exoribonucleolytically from the 5’ end, or by endoribonucleases such as the double strand-specific RNase III or the single strand-specific RNase Y, a functional analog of *E. coli* RNase E (Condon and Bechhofer 2011; Durand et al. 2012; Laalami et al. 2021; Lehnik-Habrink et al. 2011; Trinquier et al. 2020). In these pathways, ribosomes generally protect mRNAs, either by blocking the progression of RNase J1/J2 (Braun et al. 2017; Daou-Chabo et al. 2009; Mathy et al. 2007) or by masking endoribonucleolytic cleavage sites located within ORFs. In contrast to these well-described pathways that are normally inhibited by translation, we have recently identified a new RNase called Rae1 (ribosome associated endoribonuclease 1) that actually requires translation to cleave mRNA (Lalaouna and Massé 2017; Leroy et al. 2017). We solved the crystal structure of Rae1 and modeled it into the A-site of the *B. subtilis* ribosome such that it is positioned to cleave the mRNA between the A-site and P-site codons (Condon et al. 2018). Despite the predicted similarity in mechanisms, Rae1 does not share any structural homology with RelE, a member of the type II toxin-antitoxin (TA) family that binds the ribosomal A-site and typically cleaves mRNAs after the second nucleotide of the A-site codon (Hwang and Buskirk 2017), or to yeast Cue2, involved in No Go Decay (NGD), that has also been proposed to act at the A-site (D’Orazio et al. 2019).

We previously showed that Rae1 cleaves the *yrzI* operon mRNA in a translation- and reading frame-dependent manner within a short ORF called *S1025* that encodes a 17-aa peptide of unknown function (Leroy et al. 2017). The *bmrBCD* operon mRNA, encoding a heterodimeric ABC multidrug exporter formerly known as YheHI (Galián et al. 2011; Torres et al. 2009), was predicted as a second direct target of Rae1 in this global study (Leroy et al. 2017). Its expression pattern is highly similar to that of *yrzI* (correlation coefficient 0.7) in tiling array experiments performed in >100 different growth conditions in *B. subtilis* (Nicolas et al. 2012), suggesting that these two operons have linked functions. Both mRNAs were among the six most highly up-regulated mRNAs in the presence of sub-inhibitory doses of the protein synthesis inhibitor chloramphenicol (Cm) (Lin et al. 2005). Expression of the *bmrCD* portion of the operon is also up-regulated in the presence of other antibiotics that target the ribosome, including erythromycin and lincomycin, *via* a transcription attenuation mechanism thought to be triggered by antibiotic-induced ribosome pausing within the *bmrB* upstream ORF (Reilman et al. 2014).

Here, we examine the role of Rae1 in the degradation of the polycistronic *bmrBCD* mRNA and map a Rae1 cleavage site just downstream of the attenuator structure of the operon, within a newly identified short ORF, we call *bmrX*. Consistent with our previous results, we show that Rae1 also cleaves this mRNA in a translation and reading frame-dependent manner. As these two mRNAs are among the most highly expressed in the presence of low-doses of Cm (Lin et al. 2005), we asked whether this effect was Rae1-dependent and showed that over-expression of Rae1 mitigates the induction of the expression of these mRNAs by Cm. Although the absence of Rae1 renders *B. subtilis* more resistant to Cm, this phenotype is curiously not linked to the stabilization of either the *bmr* and *yrzI* mRNAs. We show that mRNA cleavage by Rae1 activates the tmRNA-dependent pathway for ribosome rescue. Overall, our results point to a role for Rae1 in mRNA quality control by eliminating mRNAs experiencing problems with translation.

## RESULTS

### The polycistronic *bmrBCD* mRNA is a direct target of Rae1

The polycistronic *bmrBCD* mRNA (Fig. 1A) was identified as a second potential target by RNAseq analysis in strains either lacking Rae1 or complemented with a plasmid over-expressing it (Leroy et al. 2017). Because the *bmrBCD* operon is regulated by AbrB at the transcriptional level (Reilman et al. 2014), we first asked whether the modulation of *bmrBCD* mRNA levels might result from an indirect impact of Rae1 on *abrB* expression. Since AbrB-mediated repression is progressively reduced in late exponential growth phase, consistent with its physiological role in transition phase phenomena (Banse et al. 2008), we performed Northern blots on total RNA isolated at both mid-log exponential and stationary growth phases. As expected, *abrB* mRNA levels were higher in mid log exponential phase than in stationary phase (Fig. 1B). However, neither *abrB* mRNA levels nor stability (measured at times after inhibition of transcription initiation by rifampicin), were altered in the *Δrae1* strain compared to wild-type (WT) (Fig. 1B and C), indicating that Rae1 does not target the *abrB* mRNA. In contrast, *bmr* mRNA levels increased in both *Δrae1* and *ΔabrB* cells, and even further in the *Δrae1 abrB* double mutant, indicating that the expression of the *bmr* operon is modulated independently by AbrB and Rae1 (Fig. 1B). The rifampicin experiment further showed that the *bmr* mRNA was highly stabilized in the *Δrae1* strain compared to the WT, where it was barely visible at the initial time point and disappeared rapidly thereafter (Fig. 1C). The *bmr* mRNA was destabilized in a *Δrae1* strain ectopically expressing WT Rae1, but not a D7N D81N catalytic mutant, indicating that its destabilization requires Rae1 catalytic activity (Fig. 1D). These data clearly indicate that Rae1 modulates *bmr operon* expression at a post-transcriptional level by accelerating its rate of decay.

**Fig. 1.**
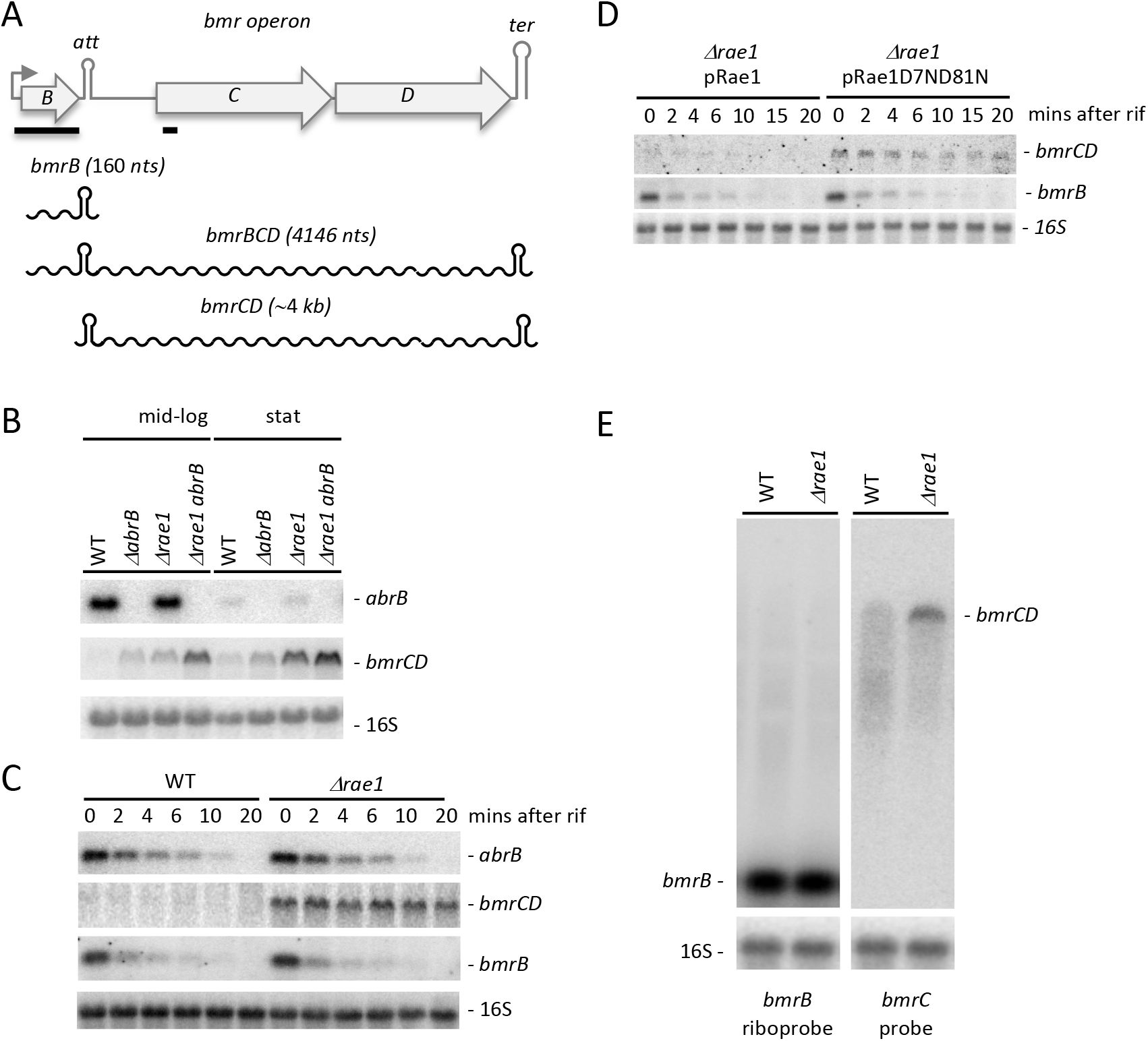
The *bmr* operon is regulated at the post-transcriptional level by Rae1. (A) Annotated structure of the *bmr* operon. ORFs are represented by grey boxes, the promoter by a rightward-pointing arrow and the transcription attentuator (att) and terminator (ter) by hairpin structures. The two primary transcripts (*bmrB* and *bmBCD*) and the processed mRNA (*bmrCD*) are shown as wavy lines and their approximate sizes are indicated. The position of the probes used is indicated by black bars. (B) Northern blot showing expression of the *bmr* and *abrB* transcripts in WT, *ΔabrB, Δrae1* and *Δrae1 ΔabrB* strains at mid-log (OD_600_ = 0.5) or stationary phase (stat) (OD_600_ = 3). (C) Northern blot showing levels of *bmr* and *abrB* mRNAs at times after rifampicin addition in WT and *Δrae1* strains at mid-log phase. (D) Northern blot showing levels of *bmr* mRNAs at times after rifampicin addition to the *Δrae1* strain containing either a plasmid expressing WT Rae1 (pRae1) or the catalytic mutant (pRae1D7ND81N) at mid-log phase. (E) Northern blot showing expression of *bmr* transcripts in WT and *Δrae1* strains at stationary phase (OD_600_ = 3). Blots were probed in a sequential order with oligo CC2344 (*bmrC*) (B-E), oligo CC2145 (*abrB*) (B and C), a riboprobe against *bmrB* (D and E), and finally with a probe complementary to 16S rRNA (CC058) as a loading control (B-E).

### Rae1 cleaves in the intergenic region between *bmrB* and *bmrC*

Since we had previously shown that Rae1 cleaves within the short *S1025* coding sequence (Leroy et al. 2017), we initially hypothesized that the Rae1 cleavage site might be located within the short *bmrB* ORF involved in antibiotic-mediated attenuation control (Reilman et al. 2014). Antibiotic-induced ribosome stalling within the short *bmrB* ORF is thought to induce formation of an anti-terminator structure and promote transcriptional read-through into the *bmrCD* structural genes to allow expression of the efflux pump. In the absence of antibiotics, transcription attenuation results in the production of a 160-nt transcript containing only *bmrB*, whereas in the presence of antibiotics, an intact readthrough *bmrBCD* transcript would be predicted to be 4146 nts in length (Fig. 1A). The ability of Rae1 to alter the stability of the short *bmrB* transcript and/or the full-length *bmrBCD* polycistronic mRNAs was examined by Northern blot analysis. The stability of the 160-nt *bmrB* transcript was similar in both the WT and the *Δrae1* strain (Fig. 1C), as well as in a *Δrae1* strain ectopically expressing WT Rae1or a D7N D81N catalytic mutant (Fig. 1D), indicating that Rae1 does not cleave the attenuated *bmrB* transcript. Strikingly, we did not detect the full-length polycistronic *bmrBCD* mRNA with a riboprobe complementary to the *bmrB* transcript (Fig. 1E), suggesting that it is rapidly processed to a shorter form containing only *bmrC* and *bmrD*, and further suggesting that the Rae1 cleavage site is located downstream of *bmrB*, within the ~4 kb *bmrCD* portion of the transcript (Fig. 1A). To narrow down the location of the Rae1 cleavage site, we generated a series of plasmids containing the *bmrC* ORF with different truncations from the 5’ end (Fig. 2A). The p*Pspac-bmrBC* construct begins 5 nts downstream of the native transcription start site, p*Pspac-hp-bmrC* starts 9 nts upstream of the attenuator and p*Pspac-bmrC* starts 40 nts upstream of the *bmrC* start codon (Fig. 2A). All constructs were placed under the control of the IPTG-inducible *Pspac* promoter to homogenize the 5’ ends of the resulting mRNAs and to avoid potential transcriptional regulation by AbrB. The Rae1 sensitivity of the different mRNAs transcribed from these plasmids was analyzed by Northern blot by comparing their steady state levels in WT and *Δrae1* strains. The mRNAs from the p*Pspac-bmrBC* and p*Pspac-hp-bmrC* constructs both accumulated in the *Δrae1* strain, while that from the shorter p*Pspac-bmrC* construct did not (Fig. 2B), suggesting that the Rae1 cleavage site is located between the attenuator and the start of *bmrC*. The lower amounts of RNA transcribed from the p*Pspac-hp-bmrC* construct are likely to be due to an inhibitory effect of a hairpin structure close to the transcription start site.

**Fig. 2.**
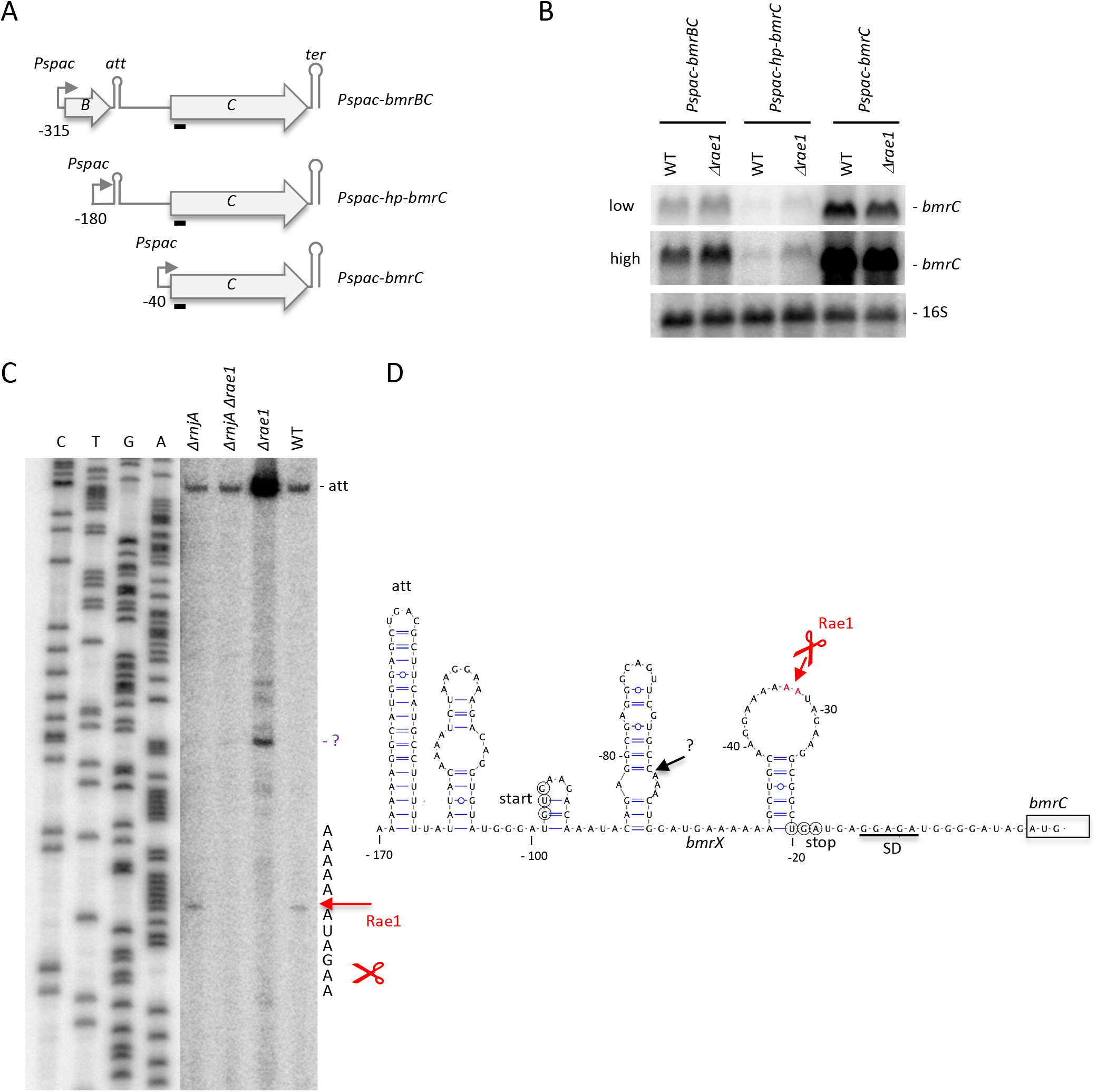
The Rae1 cleavage site is located within the intergenic region between *bmrB* and *bmrC*. (A) Structure of the p*Pspac-bmrBC*, p*Pspac-hp-bmrC* and p*Pspac-bmrC* plasmids. ORFs are represented by grey boxes, the promoter by a rightward-pointing arrow and the transcription terminator by a hairpin structure. Co-ordinates are given relative to the AUG start codon of *bmrC* Northern blot showing expression of the plasmid-derived transcripts in WT and *Δrae1* strains containing either the p*Pspac-bmrBC*, p*Pspac-hp-bmrC* or p*Pspac-bmrC* plasmid at mid-log phase (OD_600_ = 0.5). Blots were probed with oligo CC2344 (*bmrC)* (black bar in panel A) and then with a probe complementary to 16S rRNA (CC058) as a loading control. (C) Primer extension assay (oligo CC2344 within *bmrC*) on total RNA isolated from wild-type and strains lacking Rae1 (*Δrae1*), RNase J1 (*ΔrnjA*) and both (*Δrae1 ΔrnjA*) at stationary phase (OD_600_ = 3). Sequence lanes are labelled as their reverse complement to facilitate direct read-out. The mapped 5’ ends are shown to the right of the autoradiogram. A reverse transcriptase (RT) stop corresponding to attenuator (att) is indicated. This experiment was repeated twice. (D) Predicted secondary structure (Mulfold: http://www.unafold.org/mfold/applications/rna-folding-form.php) of the *bmrB-bmrC* intergenic region. Coordinates are given relative to the AUG start codon of *bmrC* (boxed). The two 5’ extremities mapped by RT are shown. The predicted attenuator stem loop (att) is shown and the Shine Dalgarno (SD) sequence of *B. subtilis* 16S rRNA is underlined. The start and the stop codon of the predicted *bmrX* ORF are circled.

To map the Rae1 cleavage site precisely, we performed primer extension assays on total RNA isolated from WT and *Δrae1* strains, using an oligonucleotide hybridizing to the early *bmrC* coding sequence. We also performed this experiment in a *ΔrnjA* background to protect the downstream cleavage product from 5’-3’ degradation, if necessary. A reverse transcriptase (RT) stop at the base of the attenuator structure was detected in all strains, and this band was stronger in the *Δrae1* mutant (Fig. 2C), consistent with the higher levels of the *bmrCD* mRNA in this strain. In the WT and *ΔrnjA* strains, we observed an RT stop mapping to 32 nts upstream of the *bmrC* start codon that was absent in both the *Δrae1* and *Δrae1 rnjA* double mutant (Fig. 2C), suggesting that this is the site of Rae1 cleavage. In the *Δrae1* strain, we detected an additional strong RT stop 30 nts upstream of the Rae1 site (Fig. 2C). It is not clear whether this stop is caused by secondary structure in this location or whether another enzyme cleaves at this site in absence of efficient degradation initiated by Rae1. The positions of the mapped 5’ ends are shown on the predicted secondary structure of the *bmrB-bmrC* intergenic region (Fig. 2D).

### Rae 1 cleaves the polycistronic *bmrBCD* mRNA within a newly identified small ORF

We had previously shown that Rae1 cleaves within the *S1025* ORF in a translation-dependent manner. We thus searched for a possible unannotated ORF in the *bmrB-C* intergenic region that overlapped the mapped Rae1 cleavage site. We anticipated that this new ORF would be truncated in the p*Pspac-bmrC* (−40) construct to explain why it was insensitive to Rae1 despite containing the cleavage site (Fig. 2B and 2D). We first made plasmid fusions of the native *bmr* promoter (*Pbmr*) to the first 301 nts of the *bmrBCD* operon including the 30 nts upstream of the *bmrC* start codon to GFP, in all three reading frames (Fig. 3A). These three constructs were introduced into WT strain and assayed for GFP production directly on agar plates. Only reading frame 0 promoted GFP expression (Fig. 3B, left), corresponding to a potential 26-aa ORF that we named *bmrX* (Fig. 3C). Previously published ribosome profiling data was consistent with ribosome binding to the predicted GUG start codon of *bmrX* (Li et al. 2012) (Fig. S1). Reinforcing this data, mutation of the GUG to a stop codon (UAG) abolished GFP expression (Fig. 3B, right). Rae1 cleavage occurred precisely between an AAA (Lys) and AUA (Ile) codon at positions 22 and 23 of the *bmrX* ORF (Fig. 2D and 3C). The BmrX peptide is conserved in several other *Bacillus* species, in particular the last four amino-acids, with the Lys-Ile codon context of the Rae1 cleavage site conserved in a subset of these peptides (Fig. 3D).

**Fig. 3.**
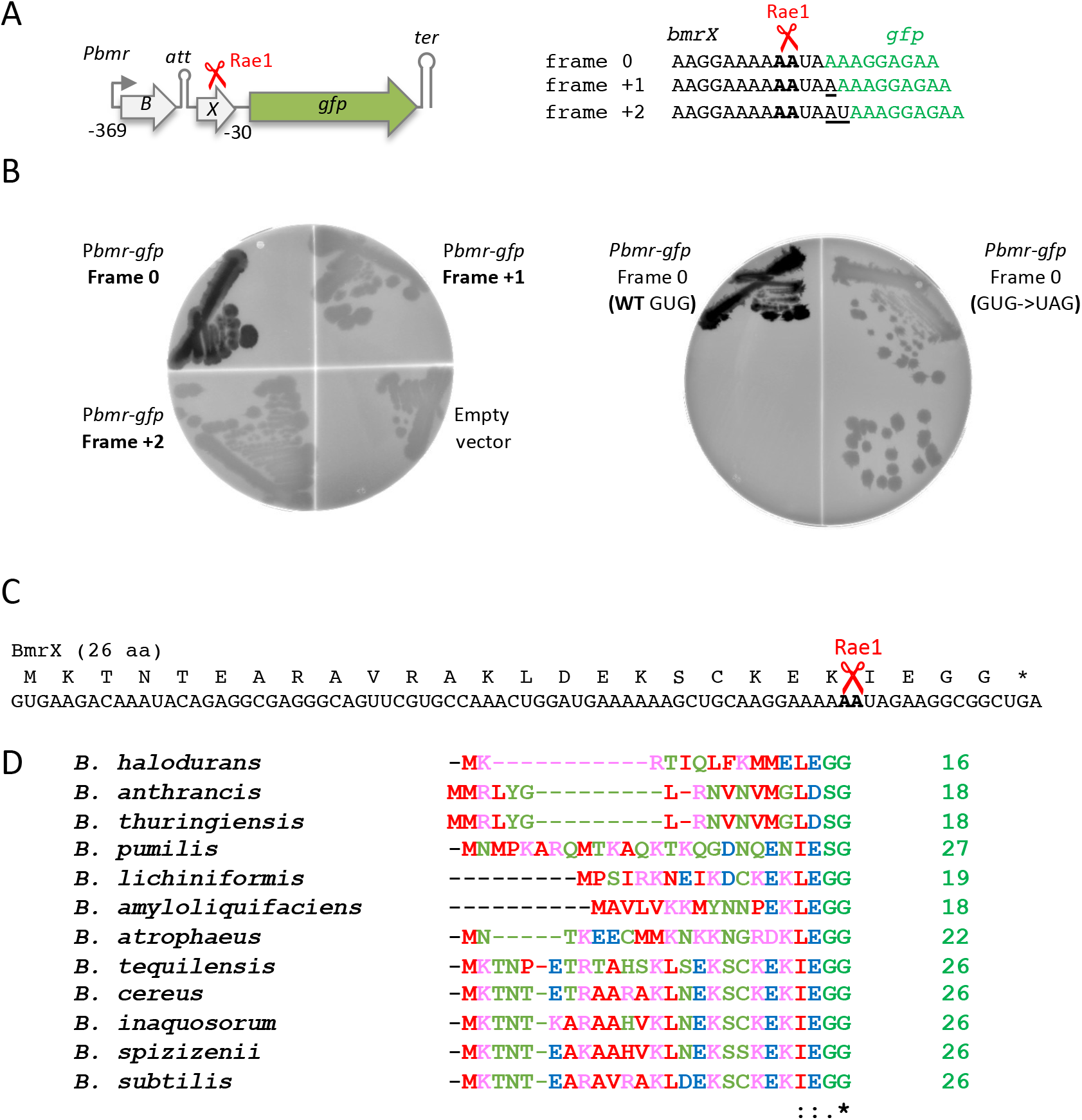
Rae1 cleaves within a newly identified small open reading frame, *bmrX*, located between *bmrB* and *bmrC*. (A) Structure of the p*bmrBXgfp* plasmid. ORFs are represented by colored boxes, the promoter by a rightward-pointing arrow, the attenuator (att) and transcription terminator (ter) by hairpin structures and the Rae1 site by a scissors symbol. Co-ordinates are given relative to the AUG start codon of *bmrC*. The sequence of the junction of the *gfp* gene with the *bmrBC* intergenic region in the 3 frames is shown. The *gfp* sequence is indicated in green. Rae1 cleaves between the two A-residues indicated in bold. (B) Fluorescence of WT strains expressing GFP from the fusion constructs p*bmrBXgfp* in the three different frames (0, +1 or +2) and of the WT containing an empty vector (left panel). Fluorescence of WT strains expressing GFP in frame 0 containing either the GUG start codon or the UAG mutant (right panel). (C) Nucleotide and amino acid sequence of *bmrX*. The localization of the Rae1 cleavage site is represented by a scissors symbol and nucleotides are in bold. (D) Conservation of the BmrX amino acid sequence in Bacilli. The sizes of the peptide encoded are indicated. (*), (:) and (.) indicate identical amino acid residues, conserved substitution, and semi-conserved substitutions respectively.

We had previously shown that insertion of a fragment containing the *S1025* ORF into the 3’UTR of the highly stable *hbsΔ* reporter mRNA rendered this mRNA sensitive to Rae1 (Leroy et al. 2017). We therefore asked whether exchanging *S1025* in the *hbsΔ-S1025* construct with the *bmrX* ORF (Fig. 4A) would promote a similar destabilization of the transcript. The *hbsΔ-bmrX* construct yields two primary transcripts from promoters P3 (P3-ter) and P1 (P1-ter) and a highly stable ribosome-protected species (R-ter). As was observed for *hbsΔ-S1025*, all three *hbsΔ-bmrX* transcripts showed increased stability in the *Δrae1* strain compared to WT. Indeed, the R-ter species was fully stabilized in the absence of Rae1, while only traces of it were detected in WT cells (Fig. 4B) showing that, like *S1025*, the *bmrX ORF* can sensitize the *hbsΔ* mRNA to Rae1. To further corroborate this result, we made additional *S1025* and *bmrX* constructs without the *hbsΔ* moiety expressed from a constitutive variant of the *Pspac* promoter lacking the *lac* operator (*Pspac (con)*). The steady state levels of the short *S1025* transcript were highly increased in the Δ*rae1* strain compared to WT (Fig. 4C), as observed with the *hbsΔ-S1025* construct (above) and the endogenous polycistronic transcript (Leroy et al. 2017). An accumulation of the short *bmrX*-containing transcript was also observed in the absence of Rae1, confirming that the presence of the *bmrX* ORF also sensitizes this mRNA to Rae1, albeit to a lesser degree than *S1025* (Fig. 4C). These data confirm that the *bmrX* ORF contains a *bona fide* and transposable Rae1 cleavage site.

**Fig. 4.**
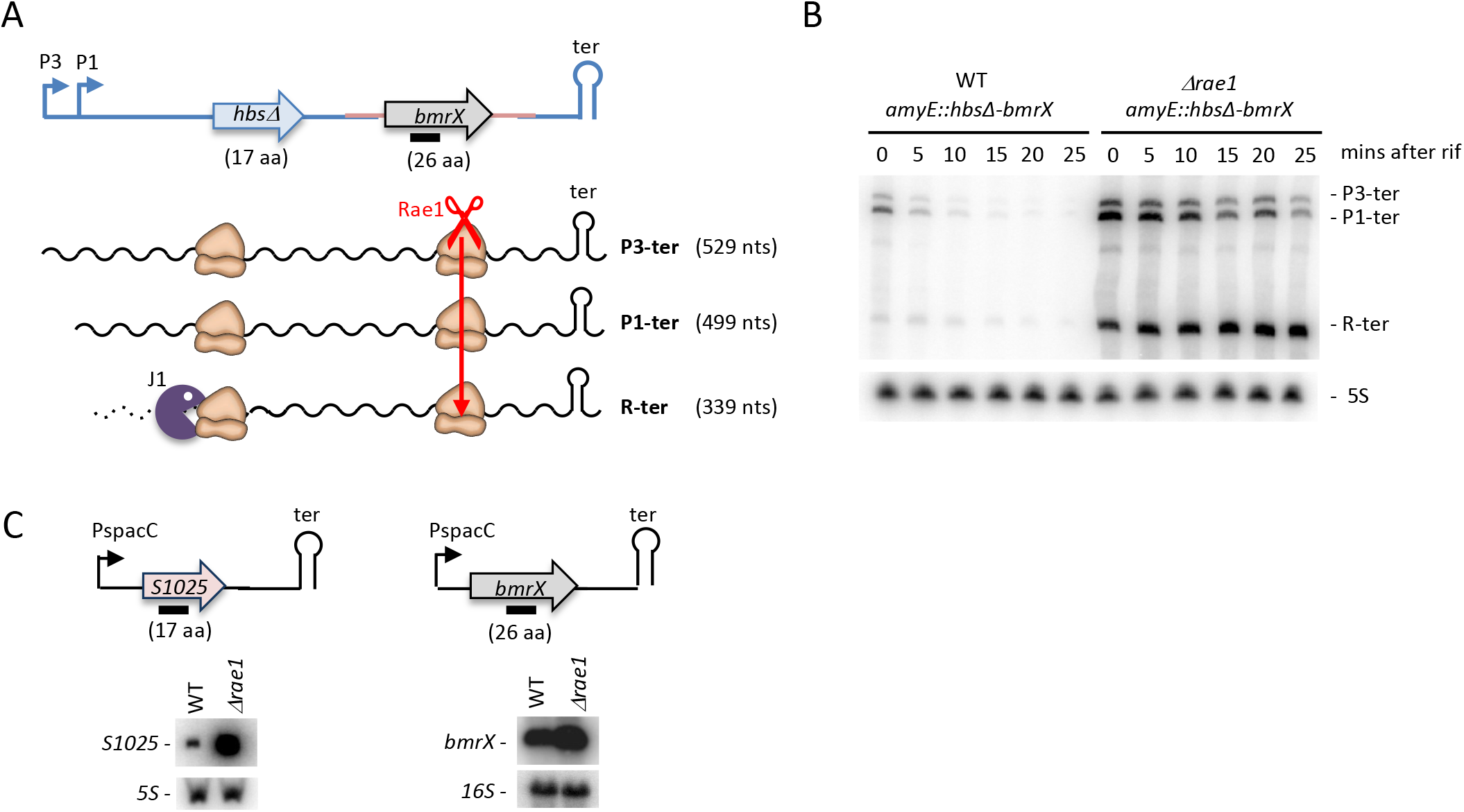
The *bmrX* ORF contains a *bona fide* and transposable Rae1 cleavage site. (A) Schematic of the two primary *hbsΔ* transcripts (P3-ter and P1-ter) and of the ribosome protected species (R-ter; 339 nts). The *hbsΔ* sequence is in blue, the 5’ -UTR and 3’-UTR of S1025 in red and the *bmrX* ORF in grey. Rae1 is depicted as a scissors symbol. The position of the probe used is indicated by a black bar. (B) Northern blot analysis of WT and *Δrae1* strains harboring the *hbsΔ-bmrX* construct in the *amyE* locus at times after rifampicin addition. The blot was probed with an oligo complementary to the *bmrX* ORF (CC2194; black bar in panel A). The origin of the 3 transcripts is shown to the right of the autoradiogram. Blots were rehybridised with a probe complementary to 5S rRNA (HP246) as a loading control. (C) Schematic of the *S1025* and *bmrX* constructs expressed from a constitutive *Pspac* promoter lacking the *lac* operator (*Pspac (con)*). The position of the probe used is indicated by a black bar. Northern blot analysis of WT and *Δrae1* strains containing the P*spac-S1025* or the P*spac-bmrX* plasmid probed with oligo CC2799 (*S1025*) or CC2194 (*bmrX)* and then with a probe complementary to 5S (HP246) or 16S rRNA (CC058) as a loading control.

### Cleavage by Rae1 within the *bmrX* ORF is translation-dependent

To ask whether Rae1 cleaves within the *bmrX* ORF in a translation-dependent manner, we made 4 mutations in the p*Pspac-bmrBC* (−315) construct described above (Fig. 2A), which we renamed p*Pspac-bmrBXC* to account for the newly discovered ORF. The mutations consisted of (i) replacing the *bmrX* GUG start codon by an UAG stop codon (*bmrX(UAG))*, (ii) deleting two consecutive C-residues in the *bmrX* coding sequence upstream of the Rae1 cleavage site to create a premature stop codon (*bmrX(Δ2)*) before the Rae1 cleavage site or (iii) adding two nucleotides upstream of the Rae1 cleavage site to modify the reading frame *(bmrX(FS+2))* or (iv) optimizing the Shine-Dalgarno (SD) sequence and translation start site from <u>C</u>AGG<u>U</u>G<u>U</u>AUGGGAU**<u>G</u>UG** to <u>A</u>*AGG<u>A</u>GG*<u>A</u>UGGGAU**<u>A</u>UG** to provide a better match to the 3’ end of 16S rRNA *(bmrX(SD+)*) (Fig. 5A and 5B). The impact of these 4 mutations on mRNA stability was analyzed in WT and *Δrae1* strains by Northern blot (Fig. 5C). Note that, like the mRNA derived from the native *bmr* locus, the *bmrBXC* transcript was processed upstream of the attenuator, leaving only the *bmrXC* portion detectable by Northern blot with the *bmrC* probe.

**Fig. 5.**
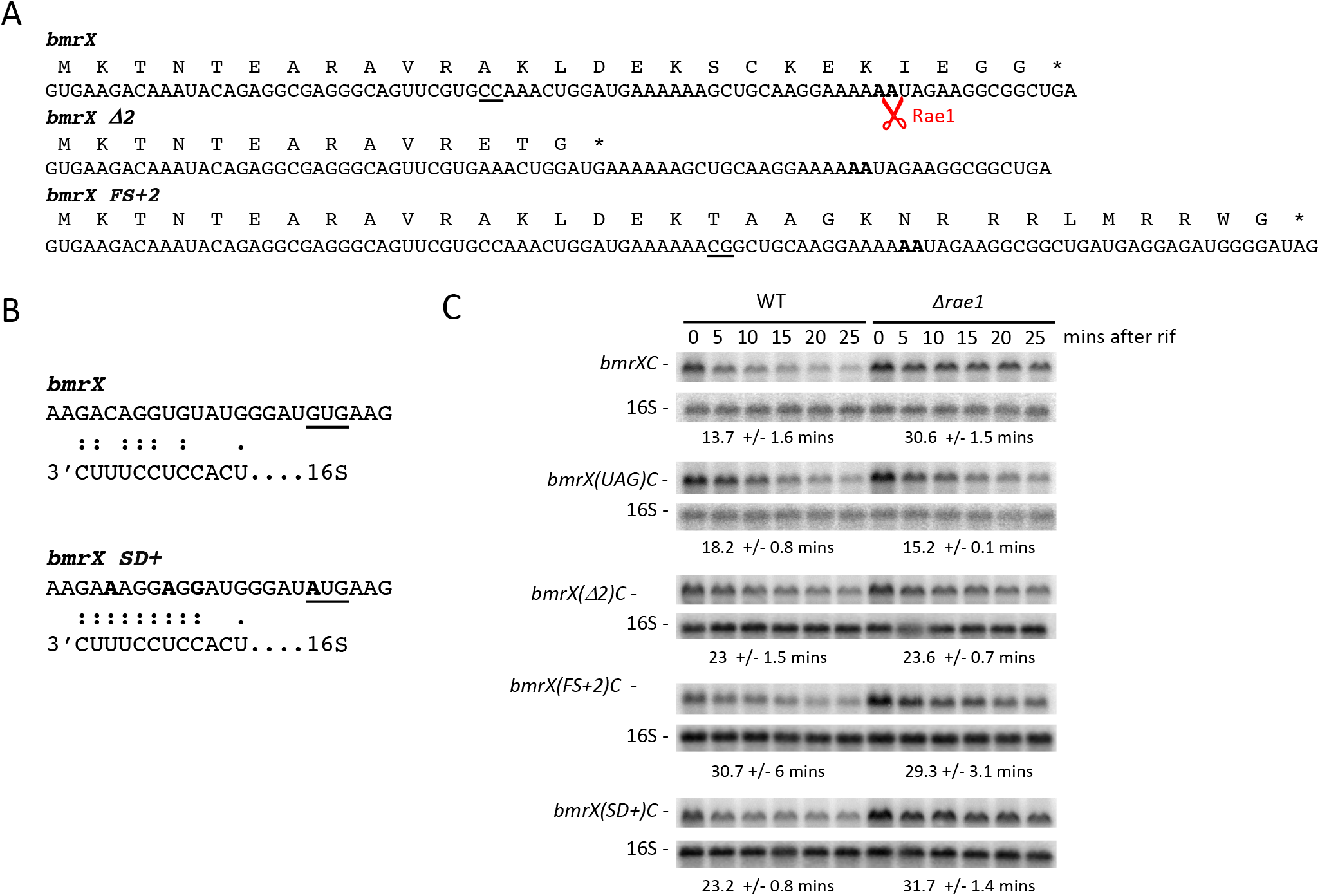
Cleavage of *bmrX* by Rae1 is translation-dependent. (A) Nucleotide and amino acid sequence of wild-type and mutant derivatives of *bmrX*. The Rae1 cleavage site is indicated in bold. The *bmrX(Δ2)* mutant contains a deletion of two C-residues (underlined in the wt *bmrX* sequence), creating a premature stop codon. In the *bmrX(UAG)* mutant, the GUG translation start codon was replaced by UAG. In the *bmrX(FS+2)* mutant, two nucleotides (underlined) were added to induce a +2 frameshift. (B) Ribosome binding sequence of the wild-type *bmrX* and the *bmrX(SD+)* mutant. The potential base-pairing with the 3′ end of 16S rRNA is indicated. The modified nucleotides in *bmrX(SD+)* mutant are in bold and the start codon is underlined. (C) Northern blot analysis of WT and *Δrae1* strains containing wt or mutant derivatives of P*spac-bmrBXC* at times after rifampicin addition, probed with oligo CC2344 (*bmrC)* and then with a probe complementary to 16S rRNA (CC058) as a loading control. Quantification of the blot presented is given in Figure S2. The calculated half-lives are the average of at least three independent experiments.

The wild-type *bmrXC* mRNA was stabilized 2.2-fold in the *Δrae1* strain (half-life of 13.7 mins in the WT vs 30.6 mins in the *Δrae1* strain) (Figs. 5C and S2). In contrast, the half-lives of the *bmrX(UAG)C, bmrX(Δ2)C* and *bmrX(FS+2)C* transcripts were all unchanged in the absence of Rae1 (Figs. 5C and S2). Thus, as previously observed for *S1025*, destabilisation of the *bmr* mRNA by Rae1 only occurs if the *bmrX* cleavage site is translated in the correct reading frame. The mRNA with the optimized translation start site, *bmrX(SD+)C*, was only stabilized about 1.4-fold (23.2 vs 31.7 mins half-life) in the absence of Rae1 (Figs. 5C and S2), suggesting that cleavage by Rae1 becomes less efficient as translation improves.

### Rae1 overexpression mitigates the induction of *S1025* and *bmrX* expression in response to chloramphenicol

A DNA microarray analysis identified the two confirmed Rae1 targets, the *bmr* and *yrzI* operon mRNAs, among the six most highly up-regulated mRNAs in *B. subtilis* following addition of sub-lethal concentrations of chloramphenicol (Cm) (Lin et al. 2005). To check whether the link between Rae1 sensitivity and induction by Cm was confined to these two operons, we first tested whether the other 4 mRNAs identified in this study (*gapB, mcpB, rbsB and ysbAB)* were also Rae1 substrates. We failed to detect the *mcpB* mRNA in either strain, even in presence of sub-lethal concentrations of Cm (data not shown), and deletion of *rae1* had only a minor effect, if any, on the stability of the *gapB, rbsB and ysbAB* mRNAs, compared to the effect on *yrzI* and *bmrCD*, suggesting that Rae1 is not a major player in the degradation of these mRNAs (Fig. S3).

To further explore the link between Rae1-sensitivity and Cm-induction of the *yrzI* and *bmr* operon mRNAs, we asked whether the increased expression of these operons in the presence of Cm was Rae1-dependent. We extracted RNA from WT and *Δrae1* cells 0, 15, 30 or 60 minutes after the addition of 0.1 or 0.5 times the minimum inhibitory concentration (MIC) of Cm (5 μg/mL).

Northern blot analysis showed that both the *yrzI* and *bmrXCD* operon mRNAs accumulated upon Cm addition to WT cells (Fig. 6A), in agreement with the published data (Lin et al. 2005). As observed previously (Leroy et al. 2017), 3 transcripts were detected for *yrzI* : the two primary transcripts P1-T4 and the P3-T4 from the P1 and P3 promoters to the main operon terminator (T4) and the matured R-T4 transcript extending from 21 nt upstream of the *yrzI* coding sequence to the T4 terminator. A very similar qualitative and quantitative pattern was observed in the absence of Rae1, suggesting that the increased expression of these operons in the presence of Cm in WT cells is Rae1-independent.

**Fig. 6.**
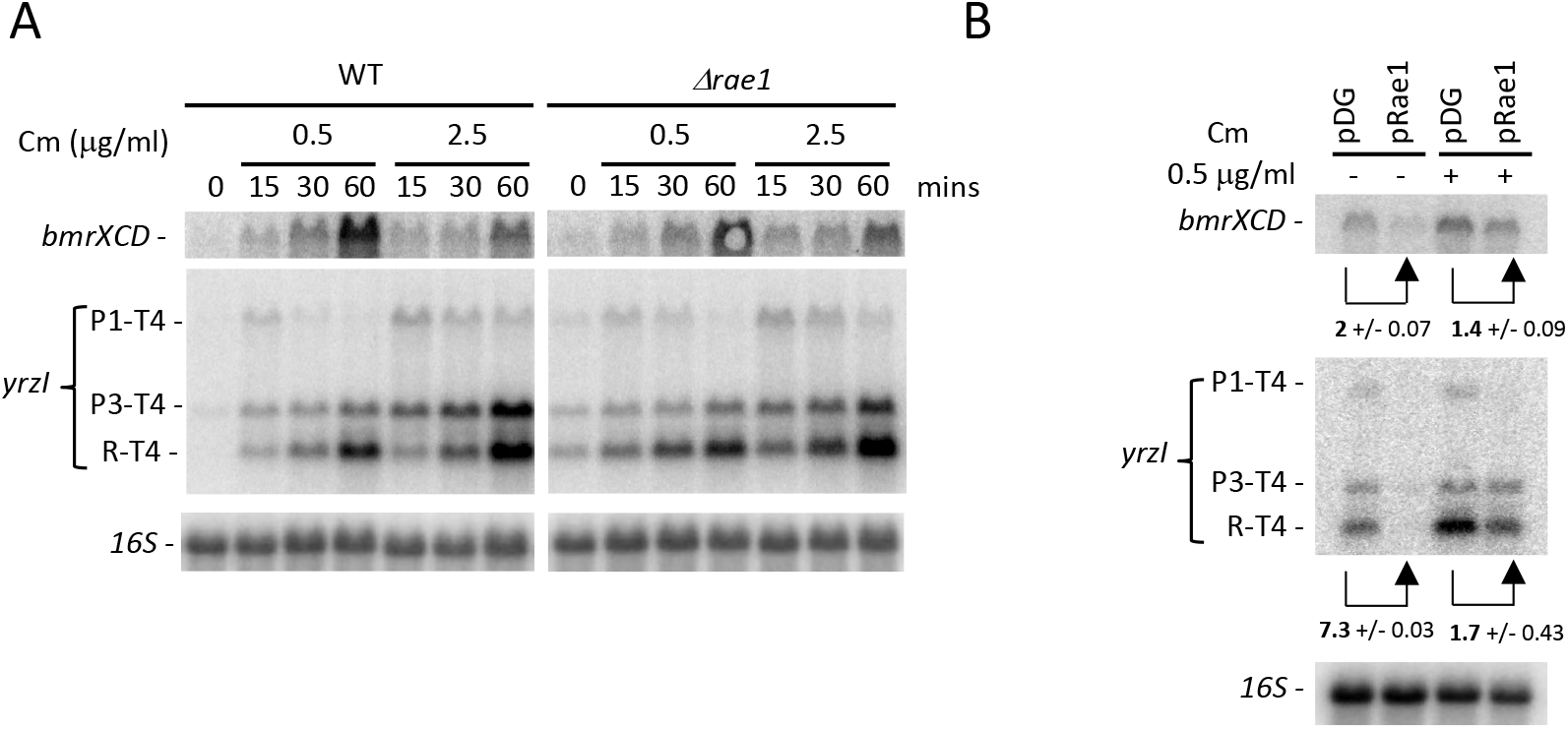
Overexpression of Rae1 mitigates the induction of *S1025* and *bmrX* expression in response to chloramphenicol. (A) Northern blot analysis of total RNA upon addition of sub-inhibitory concentrations of chloramphenicol (Cm) to exponentially growing cultures at 0.5 μg/ml and 2.5 μg/ml. Chloramphenicol was added for the times indicated. (B) Northern blot analysis of total RNA isolated from a *Δrae1* strain containing either an empty vector (pDG) or a plasmid expressing the wild-type protein Rae1 (pRae1) at mid-log phase in rich medium supplemented, or not, with 0.5 μg/ml chloramphenicol. Quantifications of the impact of Rae1 overexpression on mRNA levels from two independent experiments are indicated (A-B) Northern blots were probed in a sequential order: oligo CC2344 (*bmrC*), oligo CC1589 (*yrzI*) and then with a probe complementary to 16S rRNA (CC058) as a loading control. For *yrzI*, the correspondence between bands and transcripts is given to the right of the blot: the P1-T4 and the P3-T4 correspond to primary transcripts from the P1 and P3 promoters to the main operon terminator (T4). R-T4 refers to the 521-nt species extending from 21 nt upstream of the *yrzI* coding sequence to the T4 terminator.

We next asked the reverse question, *i.e*. whether over-expression of Rae1 could interfere with the induction of the expression of these mRNAs by Cm. We compared *yrzI* and *bmrXCD* mRNA levels in *Δrae1* cells containing either an empty vector (pDG) or a plasmid expressing the wild-type Rae1 protein (pRae1) in the presence of a sub-lethal concentration of Cm (0.1 MIC). Northern blot analysis showed that the effect of antibiotic addition was diminished for both mRNAs when Rae1 was overexpressed (Fig. 6B), suggesting that Rae1 can mitigate the induction of *yrzI* and *bmrXCD* expression in response to sub-lethal concentrations of Cm (Fig. 6B). The reduction in mRNA levels was weaker in presence of Cm than in its absence: 1.4-fold versus 2-fold for the *bmrXCD* transcript and 1.7-fold versus 7.3-fold for the *yrzI* R-T4 transcript (Fig. 6B), suggesting that Rae1 is less efficient at promoting the degradation of its targets in the presence of Cm. Whether this is due to a specific effect of Cm on Rae1 binding to the ribosome or a general effect on translation remains to be seen (it could, for example, have been anticipated that slowing translation without inhibiting it completely might actually favor Rae1 activity).

### Rae1 deletion strains show increased resistance to chloramphenicol

The stabilization of two Cm-inducible mRNAs in the *rae1* deletion strain, in particular that encoding the multidrug transporter BmrCD (Torres, 2009), led us to investigate whether this strain might show altered resistance to Cm. We performed spot dilution assays of log-phase cultures of WT and *Δrae1* strains, and *Δrae1* strains complemented with a plasmid expressing either WT or the Rae1 D7N D81N catalytic mutant. Strikingly, the *Δrae1* strain containing the empty vector was about 2-logs more resistant to 2 μg/mL Cm than the WT strain carrying the same plasmid (Fig. 7A). The *Δrae1* strain complemented with a plasmid expressing WT Rae1 (pRae1) was remarkably even more sensitive than the WT control strain to 2 μg/mL Cm, while the strain complemented with the catalytic mutant (pRae1D7ND81N) remained about 2 logs more Cm resistant (Fig. 7A).

**Fig. 7.**
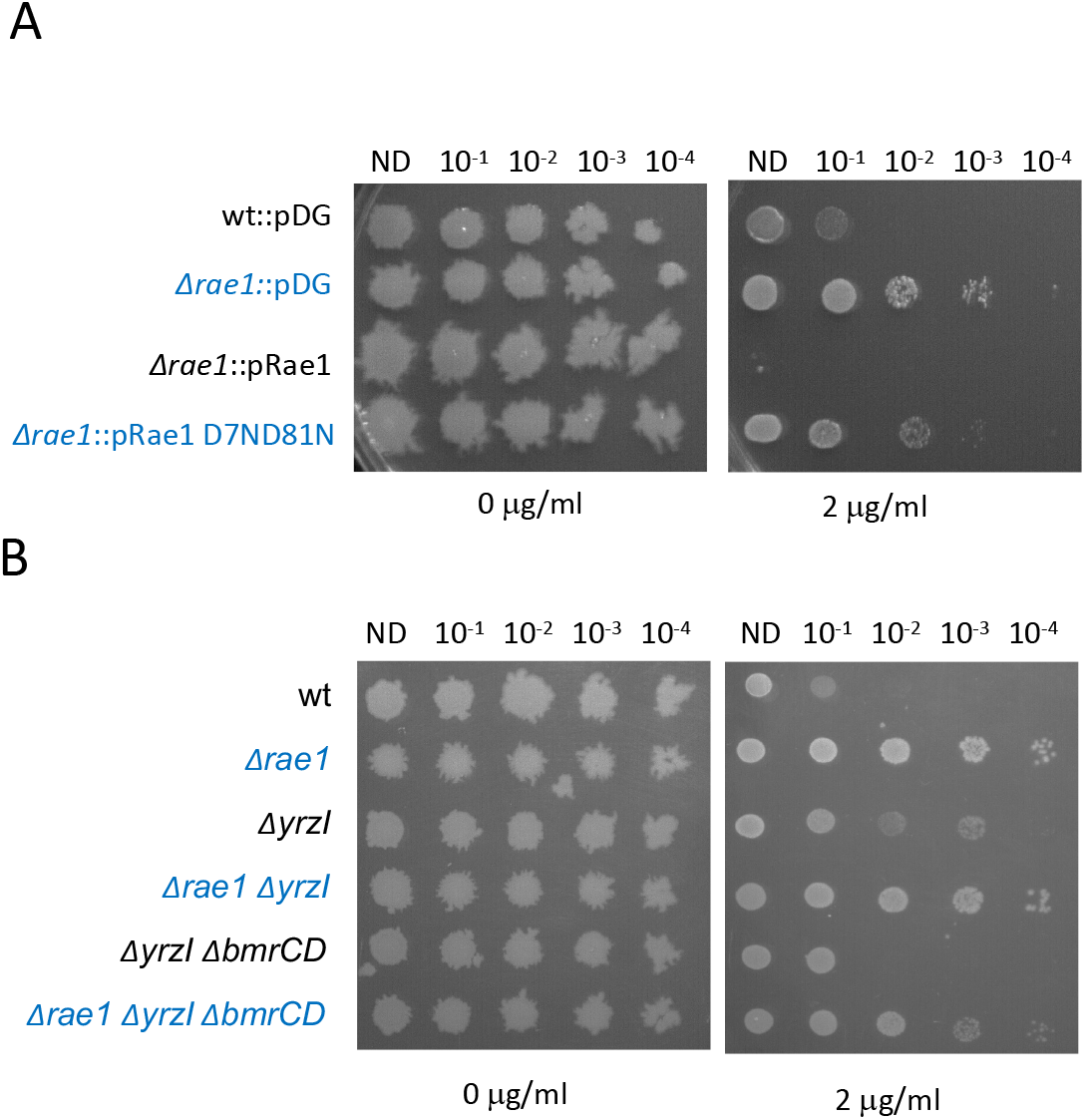
Rae1 deletion strains show increased resistance to chloramphenicol. (A) Spot dilution (10-fold) assays WT and *Δrae1* strains containing an empty vector pDG148 (pDG), and *Δrae1* strains containing a plasmid expressing either the wild-type protein (pRae1) or the catalytic mutant (pRae1D7ND81N), grown to mid-log phase (OD_600_=0.6) in rich medium, spotted on LB plates with 0 or 2 μg/mL of Cm. (A and C) IPTG (1mM) and kanamycin (5 μg/mL) were added to induce the expression of Rae1 and maintain the plasmid in bacteria, respectively. (B) Spot dilution (10-fold) assays of WT, *Δrae1*, *ΔyrzI, Δrae1 ΔyrzI, ΔyrzI ΔbmrCD* and *Δrae1 ΔyrzI ΔbmrBCD* mutant strains, grown in rich medium to mid-log phase (OD_600_=0.6), and spotted on LB plates with 0 or 2 μg/mL of Cm. (A and B) *Δrae1* strains are indicated in blue. Assays were repeated three times.

We next asked whether the increased resistance to Cm of the *Δrae1* strain was related to the stabilization of the *yrzI* or *bmr* operon mRNAs, by performing spot-dilution assays in *Δrae1* strains lacking one or both operons. Surprisingly, *Δrae1* strains lacking either *yrzI or bmrCD*, or both, still showed increased Cm resistance compared to the parental (Rae1^+^) strains (Figs. 7B and S4). Thus, the increased Cm-resistance of the *rae1* strain does not depend on either the *bmr* or *yrzI* operons and, by extension, the increased expression of these mRNAs. The key to the Cm-resistance phenotype must therefore lie with an effect of Rae1 on other (unknown) targets.

### Cleavage by Rae1 leads to ribosome rescue by the transfer-messenger RNA

Because the two Rae1 sites were mapped within the *bmrX* and *S1025* ORFs, we asked whether cleavage by Rae1 could lead to ribosome rescue by the transfer-messenger RNA (tmRNA). The *tmRNA* enters the A-site of bacterial ribosomes stalled on truncated mRNAs to promote tagging of the truncated peptide for degradation and turnover of the defective RNA (Himeno et al. 2014). To investigate the involvement of tmRNA, we used strains expressing a modified *tmRNA* (*ssrA-H6DD*) that adds a histidine tag (AGKTNSHHHHHHLDD) that escapes degradation *via* the two C-terminal D-residues that block the proteolytic pathway (Fujihara et al. 2002). This strain also contained a reporter gene, consisting of GFP fused in frame to the N-terminus of the Rae1 target S1025 (*gfp-S1025*) integrated into the *amyE* locus (Fig. 8A). We anticipated that, if the tmRNA intervenes following Rae1 cleavage, a stable His-tag would be added to the GFP-S1025 fusion protein. The *ssrA-H6DD* strain containing the *gfp-S1025* fusion was transformed with either the plasmid expressing Rae1 (pRae1) or the empty vector. We first verified that the *gfp-S1025* mRNA was sensitive to Rae1 in this background by showing that it was destabilized in the strain overexpressing Rae1 compared to the empty vector (Fig. 8B). A weak band (*) that migrates just below the full-length *gfp-S1025* transcript (940 nts) was detected in cells overexpressing Rae1 that likely corresponds to the cleaved *gfp-S1025* species (anticipated size, 774 nts). Consistent with this idea, the shorter species was not detected with a probe that hybridized to the 3’ UTR downstream the Rae1 cleavage site, indicating that it is truncated from the 3’ end (Fig. 8B). To determine whether a His-tag was added to the C-terminus of the GFP-S1025 fusion, protein extracts were absorbed on Ni^2+^-NTA agarose beads, washed and eluted with imidazole and the different fractions probed for GFP by Western blot (Fig 8C). The GFP-S1025 fusion protein was strongly enriched in the elution fraction of cells overexpressing Rae1 compared to the empty vector, indicating that the Rae1-cleaved *gfp-S1025* mRNA is rescued by the tmRNA, consistent with its proposed role in mRNA surveillance.

**Fig. 8.**
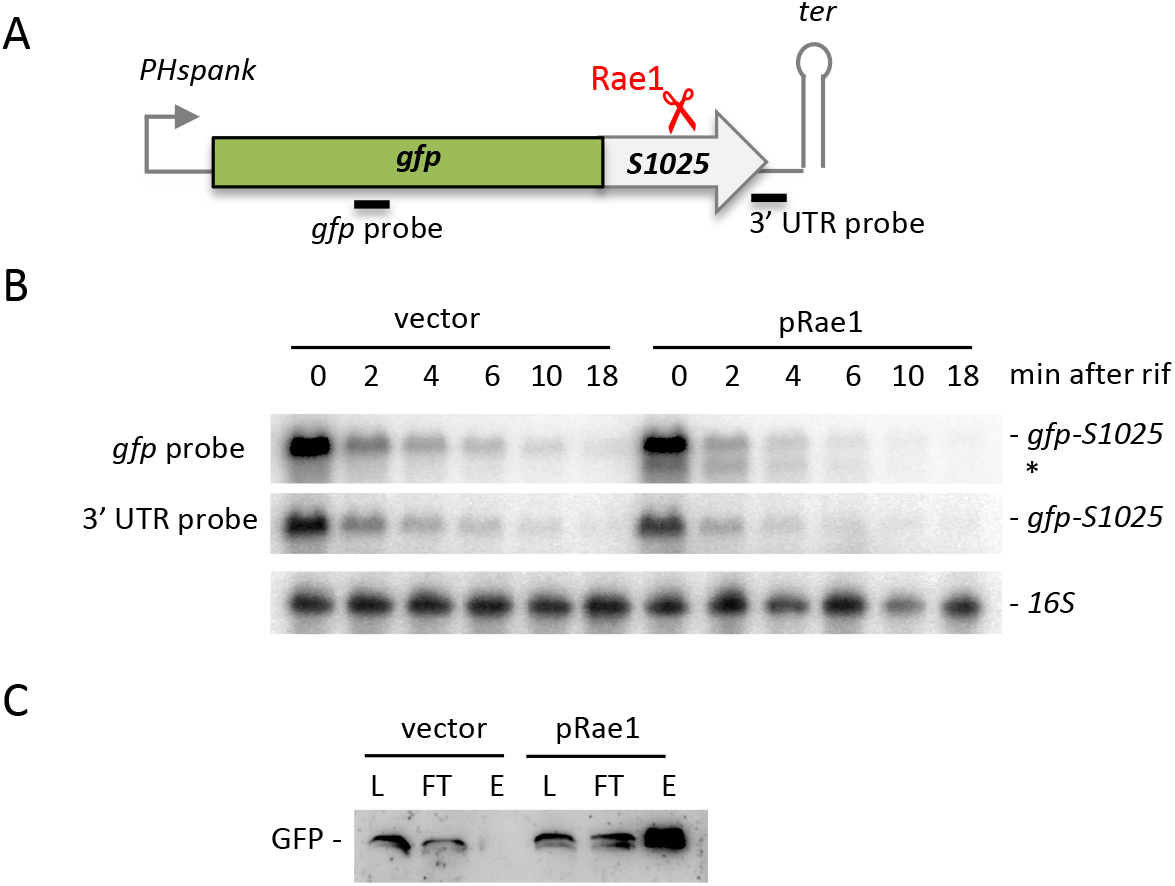
Rae1 cleavage leads to ribosome rescue by tmRNA. (A) Schematic of the *gfp-S1025* fusion. The fusion is placed under the control of the IPTG-inducible Hyperspank promoter in vector pDR111. The Rae1 site within *S1025* is indicated by a red scissors. The position of the probes used is indicated by black bars (B) Northern blot analysis at times after rifampicin addition strains containing the *gfp-S1025* fusion and mutant *ssrA* (*ssrA-H6DD*) transformed either or a plasmid expressing the wild-type protein Rae1 (pRae1) or the empty vector. The blot was probed in a sequential order with an oligo complementary to the 3’ UTR (CC429), to the *gfp* ORF (CC2422) and then with a probe complementary to 16S rRNA (CC058) as a loading control. (C) Proteins were extracted from strains used in B, absorbed, washed and eluted from Ni^2+^-NTA agarose beads. The different fractions were assayed by Western blot using GFP-specific antibodies. Fraction L corresponds to the extract loading on the beads, FT to the flow through fraction and E to the elution fraction. This experiment was repeated three times.

### MutS2 and Rae1 act in independent pathways

The recent identification of the MutS2 protein as a sensor of ribosome stalling and collisions in *B. subtilis* (Cerullo et al. 2022) lead us to wonder whether MutS2 and Rae1 work in independent pathways or cooperate to eliminate mRNAs that encounter problems with translation. MutS2 contains an SMR nuclease domain, also present in Cue2 (yeast) and SmrB (*E. coli*) (Leedom and Keiler 2022). These three proteins were shown to promote cleavage of mRNAs containing collided ribosomes, contributing to ribosome recycling (Cerullo et al. 2022; D’Orazio et al. 2019; Saito et al. 2022). To investigate a potential interplay between Rae1 and MutS2, we determined the stability of the two known Rae1 targets, *yrzI* and *bmr*, in Δ*mutS2* single and Δ*mutS2 rae1* double mutants in rifampicin experiments. Both mRNAs were unstable in WT and *mutS2* deleted strains and similarly stabilized in *rae1* and *rae1 mutS2* strains (Fig. 9), showing that Rae1 acts independently to MutS2 to cleave its targets and promote their decay. In a second assay, we compared the growth of the WT and the Δ*rae1* strain to the Δ*mutS2* and the Δ*mutS2 rae1* double mutant strains in presence of sub-lethal concentrations of Cm at 2 μg/mL in liquid cultures. In agreement with published data, the growth curves showed that the Δ*mutS2* strain was more sensitive to Cm than the WT while the *rae1* deleted strain was more resistant, consistent with our spot dilution assay data (Fig. 7A and S5). The Δ*mutS2 rae1* double mutant showed a similar level of Cm-sensitivity to the WT (Fig S5), indicating that the *rae1* and *mutS2* deletions neutralise each other’s phenotypes. Together, these results suggest that Rae1 and MutS2 act independently and use distinct mechanisms to determine *B. subtilis* Cm sensitivity levels and contribute to mRNA quality control.

**Figure 9 :**
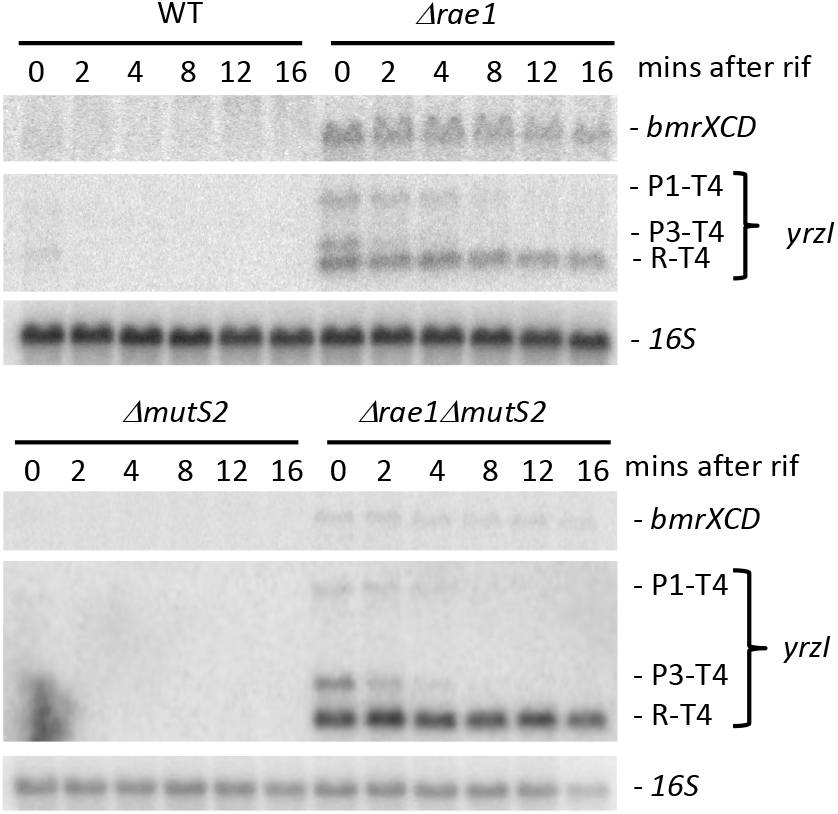
MutS2 and Rae1 act in independent pathways. Northern blot analysis of WT, *Δrae1, ΔmutS2* and *Δrae1ΔmutS2* at times after rifampicin addition. The blot was probed in a sequential order: oligo CC2344 (*bmrC*), oligo CC1589 (*yrzI*) and then with a probe complementary to 16S rRNA (CC058) as a loading control. For *yrzI*, the correspondence between bands and transcripts is given to the right of the blot: the P1-T4 and the P3-T4 correspond to primary transcripts from the P1 and P3 promoters to the main operon terminator (T4). R-T4 refers to the 521-nt species extending from 21 nt upstream of the *yrzI* coding sequence to the T4 terminator.

## DISCUSSION

We had previously mapped a Rae1 cleavage site within the small ORF *S1025* of the polycistronic *yrzI* operon mRNA (Leroy et al. 2017). To improve our knowledge on the physiological role of Rae1 and the determinants that drive Rae1 cleavage, we characterized a second target of Rae1 to determine to what extent the conclusions drawn in our previous study could be generalised. We showed that Rae1 is directly involved in the degradation of the polycistronic *bmr* multi-drug transporter mRNA (Torres et al. 2009), previously identified as a potential target by RNAseq (Leroy et al. 2017). Our unpublished data and experiments from (DeLoughery et al. 2018) suggest that the full-length *bmrBXCD* transcript is initially processed by RNase Y upstream of the attenuator hairpin, which likely protects the *bmrXCD* portion of transcript from 5’-exoribonucleolytic degradation. The processed *bmrXCD* transcript accumulates strongly in the *Δrae1* strain, indicating that Rae1 plays a key role in its turnover. Thus, *bmr* operon expression is regulated at multiple levels, including transcriptional repression by AbrB during exponential growth phase, transcriptional attenuation upon antibiotic induced ribosome stalling within the *bmrB* ORF (Reilman et al. 2014) and, as shown here, at the post-transcriptional level by Rae1 cleavage. Our data suggest that transcriptional attenuation does not efficiently block transcription of the *bmrCD* portion of the mRNA in absence of antibiotics, revealed by the strong accumulation of this transcript in the absence of Rae1. Thus, these two mechanisms together apply ‘belt and braces’ security to fully turn off the expression of this operon in the absence of substrates. In the presence of antibiotics, such as Cm, Rae1 appears to be rapidly overwhelmed by the readthrough transcript and has little impact on its accumulation (Fig. 6A), suggesting Rae1 might be present in only limiting amounts in the cell.

The Rae1 cleavage site lies between the attenuator and *bmrC*, in an unannotated small ORF, we called *bmrX*. Rae1 cleaves this ORF between a lysine codon (AAA) in position 20 and an isoleucine codon (AUA) in position 21, in a well-conserved amino acid sequence E<u>KI</u>EGG in the Bacillales (Fig. 3D). Similar to *S1025*, insertion of the *bmrX* ORF into the 3’UTR of the highly stable *hbsΔ* mRNA promotes the destabilisation of this mRNA in a Rae1-dependent fashion. These two Rae1 targets share common four features: they both encode a small peptide, both are cleaved by Rae1 when they are translated in a specific reading frame, both contain a transposable Rae1 cleavage site within their ORFs and both are upregulated in response to sub-lethal concentrations of Cm. However, there is no obvious relationship between the *bmrX* and *S1025* cleavage sites, with the latter occurring between a glutamate codon (GAG) in position 13 and a lysine codon (AAG) in position 14 in a conserved M<u>EK</u>DQV sequence (Leroy et al. 2017).

We previously hypothesized that Rae1 cleavage might occur when ribosomes pause long enough to allow Rae1 to access to mRNA (Condon et al. 2018; Leroy et al. 2017). Because the reading frames of *S1025* and *bmrX* are both critical for cleavage, if ribosome arrest does occur, it would most likely result from interaction between these nascent polypeptide chains with the ribosome exit tunnel. Although the *S1025* and *bmrX* peptides do not share any homology or obvious sequence features, such as consecutive prolines or consecutive rare codons, known to promote ribosome stalling (Huter et al. 2017; Samatova et al. 2020), studies on stalling sequences generally have highlighted the very wide variety of natural stalling motifs, making them very difficult to predict (Arenz et al. 2014; Gupta et al. 2016; van der Stel et al. 2021; Su et al. 2017; Tanner et al. 2009; Wilson et al. 2016; Woolstenhulme et al. 2013). Even stalling peptides with similar functions, *e.g*. the two well-characterized peptide leaders, MifM in *B. subtilis* (Chiba et al. 2009; Sohmen et al. 2015) and SecM in *E. coli* (Nakatogawa and Ito 2001), which promote ribosome stalling via interactions with the exit tunnel during protein export, have no detectable sequence similarity.

The two mapped Rae1 cleavages in *S1025* and *bmrX* occur within their respective ORFs generating truncated mRNAs. In bacteria, the trans-translation process, mediated by the tmRNA and its partner <u>sm</u>all <u>p</u>rotein B (SmpB), rescue ribosomes stalled on truncated mRNAs (Guyomar et al. 2021; Hayes and Keiler 2010; Himeno et al. 2014). Our data show that Rae1 cleavage leads to ribosome rescue by the tmRNA *B. subtilis*, consistent with its proposed role in mRNA surveillance of mRNAs experiencing translation problems. The MutS2 protein was also recently found to be involved in ribosome rescue in *B. subtilis* (Cerullo et al. 2022). MutS2 shares an SMR-domain with its homolog SmrB in *E. coli* (Saito et al. 2022) and with the yeast Cue2 endoribonuclease involved in No Go Decay (NGD) (D’Orazio et al. 2019). All were shown to act as ribosome collision sensors, with Cue2 proposed to act in the A-site of the colliding ribosome (D’Orazio et al. 2019), and MutS2 and SmrB acting between the stalled and colliding ribosomes (Cerullo et al. 2022; Saito et al. 2022; Leedom and Keiler 2022). Rae1 does not possess an SMR domain and has no sequence homology to MutS2, SmrB or Cue2. Moreover, multiple collisions seem to be required for optimal recruitment of these factors, but the short distance from the start codon to the cleavage site of the two Rae1 targets identified thus far, 39 nts for the *S1025* ORF and 66 nts for *bmrX*, would not allow more than 2-3 ribosomes to collide on these mRNAs. Consistent with these considerations, our data suggest that MutS2 and Rae1 act in distinct pathways and on different mRNAs: deletion of the *mutS2* gene leads to greater sensitivity to Cm while a *rae1*deletion strain is more resistant, and Rae1 targets are still stabilized in bacteria that do not express MutS2.

A global gene expression study of *B. subtilis* growing in the presence of sub-inhibitory concentrations of Cm reported that the polycistronic *yrzI* or *bmrCD* transcripts were among the most highly expressed mRNAs under these conditions (Lin et al. 2005). We confirmed that the expression of both mRNAs was induced by Cm by Northern blot. Expression levels were lower in cells overexpressing Rae1, indicating that Rae1 can counteract the induction by Cm. The reverse is also true, *i.e*. Cm can interfere with Rae1 action, since the reduction in *yrzI* and *bmr* mRNA levels upon Rae1 overproduction is stronger in the absence of antibiotic than in its presence (Fig. 6B). Because Cm was shown to bind directly to the A-site crevice on the 50S ribosomal subunit, occupying the same location as the aminoacyl moiety of an A-site tRNA (Bulkley et al. 2010), it is possible that Cm and Rae1 compete for A-site binding. Another possibility is that Cm promotes ribosome stalling at context-specific sites (Marks et al. 2016) that are not fully suited to Rae1 cleavage. The fact that *B. subtilis* strains lacking Rae1 activity are more resistant to Cm than the WT strain, independently of the expression of BmrCD efflux pump or the *yrzI* operon, suggests that resistance phenotype is associated with an effect of Rae1 on other targets. One possibility we have considered is that Rae1 recruitment to Cm-paused ribosomes could lead to the irreversible loss of some key mRNAs required for cell growth. By deleting Rae1, these mRNAs would not be degraded and translation could potentially continue, albeit more slowly. Overproduction of Rae1 would render the cells more sensitive to Cm, as we have seen.

In conclusion, we show that Rae1 cleaves the polycistronic *bmrXCD* transcript within the *bmrX* ORF in a translation and reading frame-dependent manner as reported previously for the *yrzI* operon mRNA. Our current model is that the peptide sequences of BmrX and S1025 somehow promote the stalling of the ribosome and that mRNA cleavage by Rae1 permits ribosome rescue by the tmRNA. In this way, Rae1 could contribute to translation quality control in a pathway distinct from those that involve MutS2 or its homologs (SmrB in *E. coli* and Cue2 in yeast). Although BmrX is restricted to the Bacillales, as previously reported for S1025 (Leroy et al. 2017), Rae1 is far more widely conserved among Firmicutes, the Cyanobacteria and the chloroplasts of higher plants (Condon et al. 2018). Thus, the physiological importance of Rae1 likely extends well beyond these two operons in *B. subtilis*. It also remains to be seen to what extent Rae1 can contribute to mRNA surveillance in Cyanobacteria and the chloroplast.

## MATERIALS AND METHODS

### Strains and constructs

Strains and oligonucleotides used are shown in Supplemental Tables S1and S2 respectively. The *B. subtilis* strains used in this study were derivatives of W168 (lab strain SSB1002).

Plasmids containing the *bmrC* ORF with different truncations from the 5’ end (p*Pspac-bmrBXC* (pl867), p*Pspac*-*hp-bmrC* (pl828) and p*Pspac*-*bmrC* (pl829)) were constructed by PCR amplification using the forward primer CC2617, CC2451 or CC2452 and the reverse primer CC2136, containing an *E. coli* rRNA (*rrnG*) transcription terminator to limit transcription of downstream plasmid DNA. The PCR fragments were cloned between the XbaI and SphI sites of plasmid pDG148 and placed under the IPTG-inducible *Pspac* promoter (Stragier et al. 1988).

The p*Pspac-bmrBXC* plasmid was used to construct 4 plasmids containing mutations in the *bmrX* ORF: p*bmrBX(UAG)C* (pl866), p*bmrBX(Δ2)C* (pl873), p*bmrBX(FS+2)C* (pl841) and p*bmrBX(SD+)C* (pl876). They were obtained by two-fragment overlapping PCR using the p*Pspac-bmrBXC* plasmid as template. The upstream fragment was amplified with the forward primer CC2617 and the reverse primer CC2565, CC2467, CC2774 or CC2698 and the downstream fragment with the forward primer CC2566, CC2468, CC2775 or CC2699 and the reverse primer CC2136. The overlapping fragments were reamplified using oligo pair CC2617/CC2136 and cloned between the XbaI and SphI sites of plasmid pDG148 and placed under the IPTG-inducible *Pspac* promoter (Stragier et al. 1988).

The p*bmrBXgfp* fusions containing the native *bmr* promoter (*Pbmr*) and the first 301 nts of the *bmrBXCD* operon in all three reading frames fused to GFP were obtained by two-fragment overlapping PCR. They were referred as pl845 (frame 0), pl844 (frame+1) and pl846 (frame+2). The upstream fragment was amplified with the primer pair CC2132/2419 and p*Pspac-bmrBC* as a template, and the downstream fragment using the forward primers CC2521, CC2522 or CC2523 and the reverse primer CC2421, with pHM2-*hbs*Δ∷*gfp*SDwt (pl719) as a template (Braun et al. 2017). The overlapping fragments were reamplified using oligo pair CC2132/CC2421 and cloned between the XbaI and SphI sites of plasmid pDG148. The p*bmrBXUAGgfp* fusion (pl858) containing a UAG stop codon instead of the GUG start codon was obtained by two-fragment overlapping PCR. The upstream fragment was amplified with primer pair CC2132/CC2565 and the downstream fragment with primer pair CC2566/CC2421 with p*bmrBXgfp* (pl845) as a template. The overlapping fragments were reamplified using oligo pair CC2132/CC2421 and cloned between the XbaI and SphI sites of plasmid pDG148.

The pDR-*gfpS1025* (pl875) containing the *gfp* fused to the S1025 ORF was obtained by subcloning the *gfpS1025* insert from the pHM2-*gfpS1025* (pl869) between the SalI - HindIII of the integrative the pDR111 vector. The pHM2-*gfpS1025* (pl869) plasmid was obtained by two-fragment overlapping PCR. The upstream fragment was amplified with the primer pair CC2634/2635 and p*bmrBXgfp* (pl845) as a template, and the downstream fragment using with the primer pair CC2636/CC572, pHM2-*hbsΔ-S1025* (pl707) as a template (Leroy et al). The overlapping fragments were reamplified using oligo pair CC2634/CC572 and cloned between the HindIII and BamHI site of the integrative pHM2 plasmid and placed under the control of a constitutive promoter (Pspac(con).

The pHM2-*hbsΔ-bmrX* plasmid (pl908) was obtained by three-fragment overlapping PCR. The upstream fragment was amplified with primers CC1607 and CC2893, and the downstream fragment with primers CC2894 and CC572, using pHM2-*hbsΔ-S1025* (pl707) as a template (Leroy et al). The third fragment was obtained by hybridizing the oligo pair CC2875/CC2876. The overlapping fragments were reamplified using oligo pair CC1607/CC572 and cloned between the EcoRI and BamHI site of the integrative pHM2 plasmid, placed under the control the native promoters of *hbs* gene.

The pHM2-*bmrX* (pl936) and the pHM2-*S1025* (pl925) plasmids were constructed by PCR amplification using the forward primer CC2962 and the reverse primer CC572 with pHM2-*hbsΔ-bmrX* (pl908) or pHM2-*hbsΔ-S1025* (pl707) as a template. The PCR fragments were cloned between the HindIII and BamHI site of the integrative pHM2 plasmid, placed under the control of the constitutive Pspac(con) promoter.

All plasmid constructs were verified by sequencing and transformed into the strains SSB1002 (WT) and CCB375 *(Δrae1*) (Table S1). In all the plasmid constructs using the pDG148 vector, the cloned genes were followed by an *E. coli* rRNA (*rrnG*) transcription terminator to limit transcription of downstream plasmid DNA. The integrative plasmid derived from pHM2 and from pDR111 were linearized with XbaI and SacI respectively, before transformation, for integration into the *amyE* locus of SSB1002 (WT) or CCB375 *(Δrae1*) strains (Table S1). The CCB375 *(Δrae1*) and the CCB942 (*ssrADD*) strains are resistant to erythromycin and chloramphenicol, respectively. The replicative pDG148 and the integrative pDR111 and the pHM2 vectors confer kanamycin, spectinomycin and chloramphenicol resistance, respectively.

Strain CCB748 was constructed by transferring the *rnjA∷spec* construct from strain CCB434 into CCB375 *(Δrae1*) (Table S1). Stains CCB1058 and CCB1059 were constructed by transferring the *abrB∷cm* construct from strain SSB1010 into SSB1002 (WT) and CCB375 *(Δrae1*) (Table S1). Strains CCB1611 and CCB1612 were constructed by transferring the *mutS2∷kan* construct from BKK28580 (Koo et al. 2017) strain into W168 (SSB1002) or CCB375 *(Δrae1*) (Table S1).

Strains CCB1447 and CC1448 were constructed by transforming strain CCB1444 with the pRae1 plasmid (pl660) that allows Rae1 expression (Leroy et al, 2017) or the empty pDG148 vector. Strain CCB1444 was constructed in two steps: the integrative plasmid pDR-*gfpS1025* (pl875) (Table S1) was first transformed into the WT strain SSB1002 and the resulting strain (CCB1368) was transformed with the *ssrAH6DD∷Cm* construct from strain AHMG Pspac-ssrA(H6DD) (Fujihara et al. 2002).

Strain CCB1469 (*bmr∷spec*) was constructed using a PCR fragment generated by re-amplifying three overlapping PCR fragments (oligo pair CC2039/CC2847), corresponding to the upstream region of the *bmrB* gene (oligo pair CC2039/CC2037), the downstream region of the *bmrD* gene (oligo pair CC2846/CC2847) and the spectinomycin resistance cassette from plasmid pDG1727 (oligo pair CC2694/CC2695). Strain CCB1520 (*yrzI∷kan*) was constructed using a PCR fragment generated by re-amplifying three overlapping PCR fragments (oligo pair CC2848/CC2851), corresponding to the upstream region of the *S1027* gene (oligo pair CC2848/CC2849), the downstream region of the *S1024* gene (oligo pair CC2850/CC2851), and the kanamycin resistance cassette from plasmid pDG780 (oligo pair CC011/CC012). The resulting PCR fragments were used to transform *B. subtilis* W168 (SSB1002) and the correct insertion verified by PCR. Strain CCB1387 was obtained by transformation of CCB375 (*Δrae1)* with chromosomal DNA from strain CCB1469 (*Δbmr*). Strains CCB1526, CCB1527 and CCB1528 were obtained by transformation of CCB1469 (*Δbmr*), CCB375 (*Δrae1)* and CCB1387 (*Δrae1 Δbmr)*, respectively, with chromosomal DNA from strain CCB1520 (*ΔyrzI*).

### GFP detection

GFP fluorescence of bacteria was detected on agar plates using Typhoon scanner (GE) with excitation at 470 nm and emission at 520 nm.

### RNA isolation and Northern blots

Northern blots were performed on total RNA isolated from *B. subtilis* cells growing 2xYT medium either by the glass beads/phenol method described in (Bechhofer et al. 2008) or by the RNAsnap method described in (Stead et al. 2012). Northern blots were performed as described previously (Durand et al. 2012). The *bmrB* riboprobe was transcribed *in vitro* using T7 RNA polymerase (Promega) and labeled with [α-32P]-UTP using a PCR fragment amplified with oligo pair CC2195/CC2196 as template. Northern blots were exposed to PhosphorImager screens (GE Healthcare) and the signal was obtained by scanning with a Typhoon scanner (GE Healthcare) and analyzed by Fiji (ImageJ) software.

### Primer extension assays

Primer extension assays were performed on glass bead/phenol extracted RNAs as described previously (Britton et al. 2007). Oligo CC2344 was used to map the Rae1 cleavage site within the *bmrX* ORF of the *bmr* polycistronic mRNA.

### Spot dilution assays

Cells were grown in 2xYT medium to OD_600_ = 0.6. When cells contained either the empty plasmid pDG148, the pRae1 (pl660) or pRae1D7ND81N (pl664), allowing the expression of the wild-type or the Rae1 catalytic mutant, respectively, 10-fold serial dilutions in MD medium without carbon source were spotted on LB plates containing kanamycin (2.5 μg/ml) to maintain the plasmid and IPTG (1mM) to induce the expression of the *rae1* gene. Where indicated, chloramphenicol was added at 2 μg/ml.

### Ribosome rescue assay

For proteins extractions, cells from strains CCB1447 and CC1448 were resuspended in ice-cold Buffer (20 mM Tris–HCl pH 7.5, 1 mM EDTA, 50 mM NaCl) and lysed by sonication (3 times 30”). The lysate was cleared at 16,100 g for 10 min at 4°C and protein concentration was determined by the bradford method. 400 μg of protein extracts were incubated with 50 μl of Ni-NTA agarose beads overnight, washed twice with 10 volumes of 20 mM imidazole and eluted with one volume of 250 mM imidazole. The different fractions were assayed by Western blot (ECL; GE) using GFP-specific mouse antibodies (Sigma).

### Growth curves in a microplate reader

For each strain, 2xTY medium was inoculated from exponential culture at initial OD ≈ 0.02 and 200 μl were dispensed in triplicate into a 96-well microplate. Specific wells were inoculated with medium only for background correction purposes. Absorbance readings (600 nm) were taken every 30 minutes over the 12 h time period at 37°C using a BMG LABTECH microplate reader.

## ACKNOWLEDGMENTS

We thank lab members for helpful discussion. This work was supported by funds from the CNRS and Université Paris Cité (UMR8261), the Agence Nationale de la Recherche (ARNr-QC, Labex Dynamo and Equipex Cacsice). We also thank A. Muto for the gift of the strain AHMG Pspac-ssrA(H6DD).

---

**Fig. S1.**
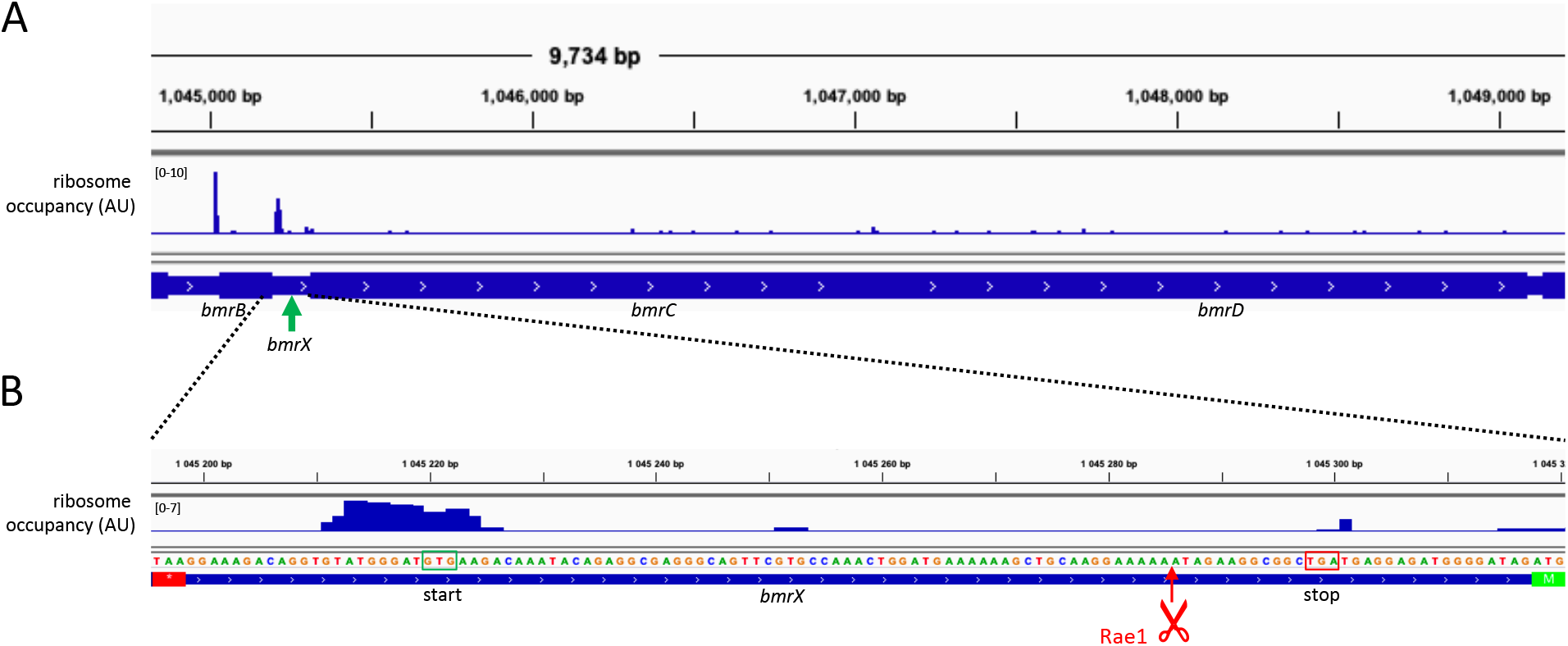
Ribosome profiling data is consistent with ribosome binding to the predicted GUG start codon of *bmrX*. (A) Ribosome occupancy profiles on the *bmrBCD* polycistronic mRNAs from Li *et al*. 2012. The location of *bmrX* is indicated by a green arrow. (B) Zoom on the *bmrX* sequence. The predicted GUG start codon and UGA stop codon of *bmrX* are boxed in green and red, respectively. The stop codon of *bmrB* and the start codon of *bmrC* are indicated as red and green bars. The Rae1 site mapped within *bmrX* is showed by a red arrow.

**Fig. S2.**
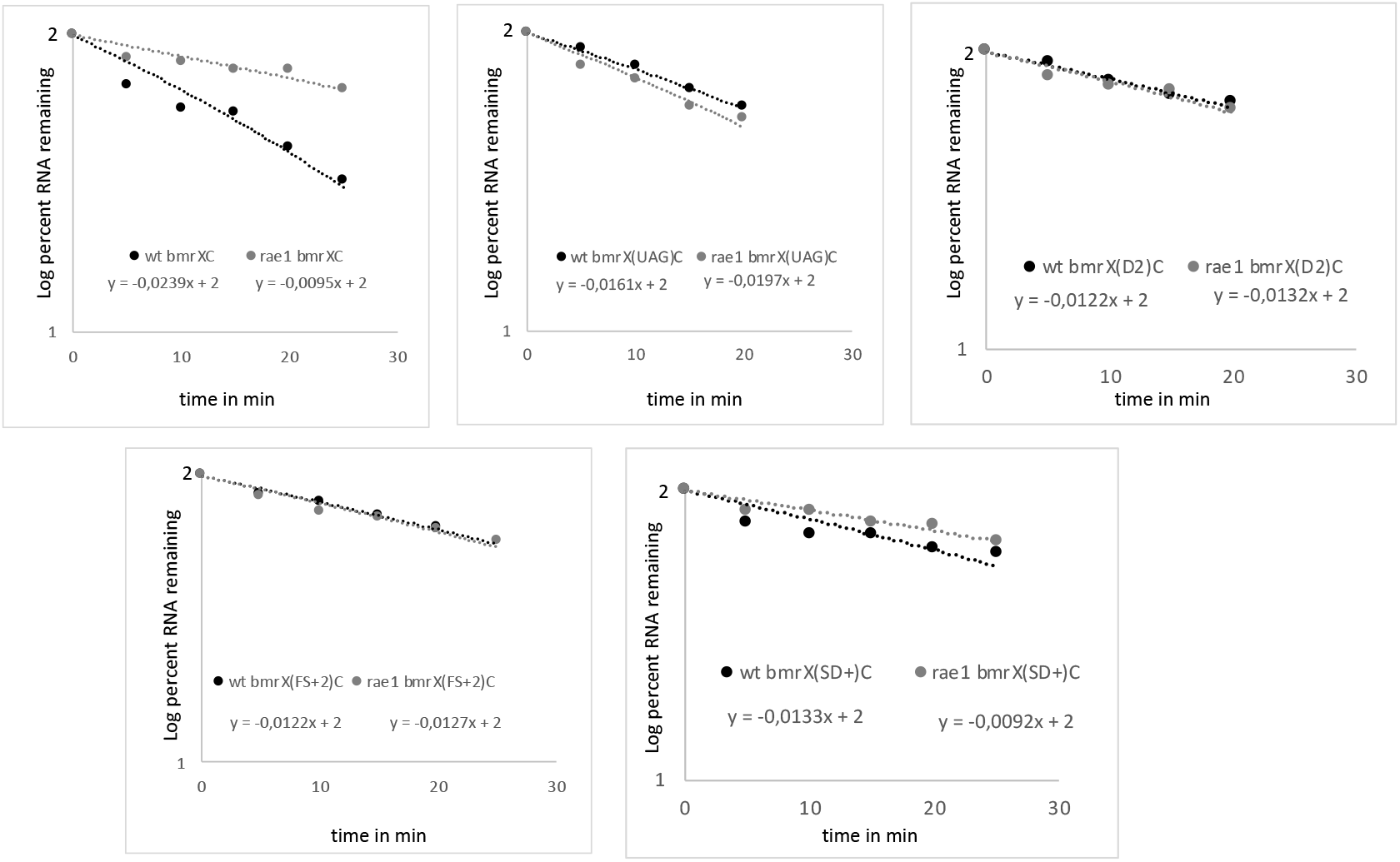
RNA decay plots of wild-type and mutant derivatives of *bmrXCD mRNAs*. WT strains are shown by black dots and *Δrae1* strains vy grey dots. The slopes of the curves used for half-life calculations are given for each plot.

**Fig. S3.**
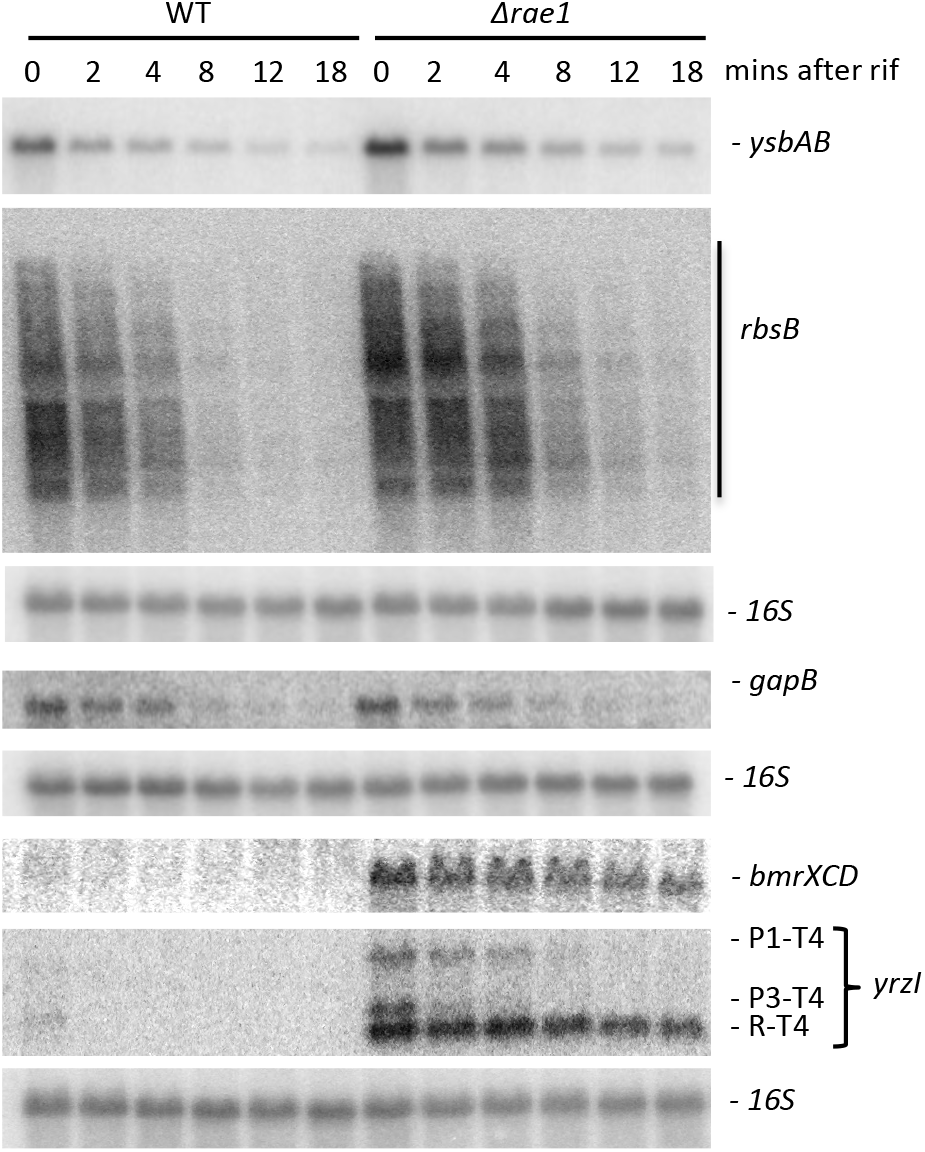
Deletion of *rae1* does not impact the stability of the *gapB*, *rbsB* and *ysbAB* mRNAs. Northern blot analysis of WT and *Δrae1* strains at times after rifampicin addition. The blot was probed with an oligo complementary to the *ysbAB* ORF (CC2405), the *rbsB* ORF (CC951), the *gapB* ORF (CC3018), the *bmrC* ORF (CC2344) or the *yrzI* ORF (1589). Blots were rehybridised with a probe complementary to 16S rRNA (CC058) as a loading control.

**Fig. S4.**
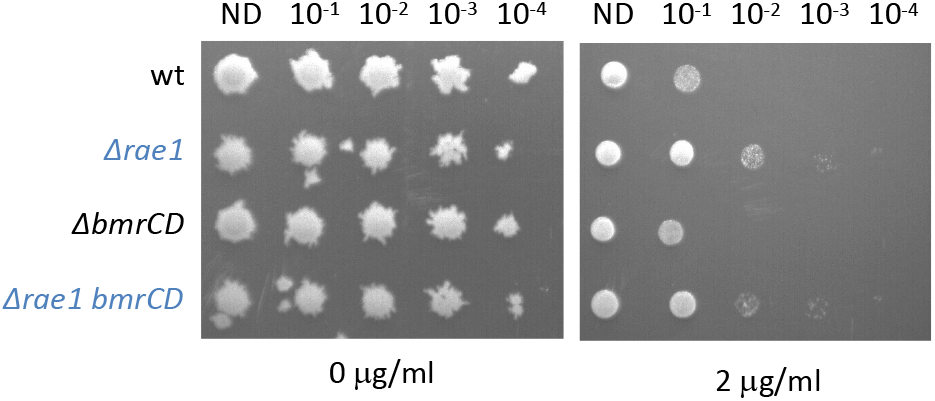
The increased Cm resistance of the *Δrae1* strain is independent of *bmrBCD*. Spot dilution (10-fold) assays of WT, *Δrae1*, *ΔbmrBCD* and *Δrae1 ΔbmrBCD* mutant strains, grown in rich medium to mid-log phase (OD600=0.6), and spotted on LB plates containing 0 or 2 μg/ml Cm.

**Fig. S5.**
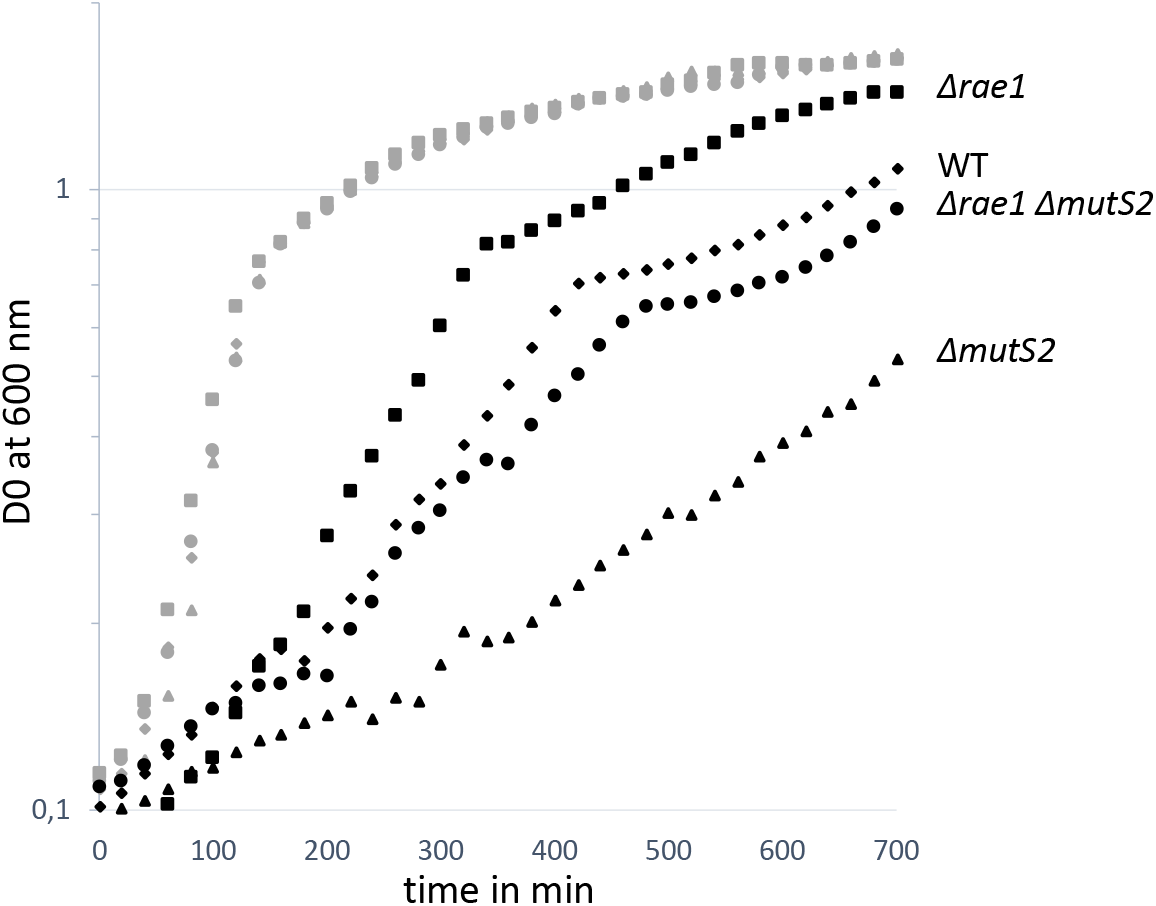
The *mutS2* and *rae1* deleted strains have different sensitivities to Cm. Growth of the WT (diamonds), the *Δrae1* (squares), the *ΔmutS2* (triangles) and *Δrae1ΔmutS2* (circles) strains at 37°C in a 96-well plate in 2xTY medium (grey symbols) or in 2xTY medium with 2 μg/ml of Cm (black symbols). OD measurements at 600 nm were performed using a BMG LABTECH microplate reader. Assays were repeated twice.

